# Electromyographic Correlates of Effortful Listening in the Vestigial Auriculomotor System

**DOI:** 10.1101/2023.07.19.549747

**Authors:** Andreas Schroeer, Farah I. Corona-Strauss, Ronny Hannemann, Steven A. Hackley, Daniel J. Strauss

## Abstract

Recently, electromyographic (EMG) signals of auricular muscles have been shown to be an indicator of spatial auditory attention in humans, based on a vestigial pinna-orienting system. Because spatial auditory attention in a competing speaker task is closely related to the more generalized concept of attentional effort in listening, the current study investigated the possibility that the EMG activity of auricular muscles could also reflect correlates of effortful listening in general. Twenty subjects were recruited. EMG signals from the left and right superior and posterior auricular muscles (SAM, PAM) were recorded while subjects attended a target podcast in a competing speaker paradigm. Three different lev-els of listening effort, low, medium, and high, were generated by varying the number and pitch of distractor streams, as well as the signal-to-noise ratio. All audio streams were either presented from a loudspeaker placed in front of the participants (0°), or in the back (180°). Averaged PAM activity was not affected by different levels of listening effort, but was significantly larger when stimuli were presented from the back, as opposed to the front. Averaged SAM activity, however, was significantly larger in the high listening effort condition, compared to low and medium, but was not affected by stimulus direction. We hypothesize that the increased SAM activity is a response of the vestigial pinna–orienting system to effortful stream segregation task.

## 1 Introduction

Recently, Strauss et al. (2020) demonstrated that electromyographic (EMG) signals of several auricular muscles, specifically the posterior, anterior, superior, and transverse auricular muscles (PAM, AAM, SAM, and TAM), are an indicator of the spatial direction of auditory attention.

This vestigial pinna-orienting system is a so-called “neural fossil” (Hackley, 2015, Strauss et al., 2020), and has a reflexive, stimulus driven component in response to transient auditory stimuli. This component has been observed as transient EMG responses by the PAM, AAM, and TAM, starting approximately 70 ms after rapid-onset auditory stimuli. This part of the vestigial pinna-orienting system does not depend on the participants’ voluntary, task-oriented focus, and is therefore referred to as exogenous attention in Strauss et al. (2020), and strongly indicates the direction of the salient auditory stimuli. The second component of this system is based on deliberately attending an audio stream, while ignoring a competing, but spatially separate stream, and is referred to as endogenous attention. In this case, Strauss et al. (2020) reported sustained activity of the PAM, AAM, and SAM, that was larger on the side of the attended audio stream that on the side of the ignored stream. Furthermore, this effect was enhanced when audio streams were presented outside of the participant’s field-of-view (*±*120°), compared to inside their field-of-view (*±*30°). Overall, Strauss et al. (2020) reported differences, especially in the SAM, between purely stimulus driven responses and responses during the active listening task in a challenging condition, that required attentional effort, see Sarter et al. (2006). Listening effort and its relation to different modes of attention and/or cognitive resource limits has been established and analyzed by several models of effortful listening, e.g., Strauss et al. (2010), McGarrigle et al. (2014), Pichora-Fuller et al. (2016), Strauss & Francis (2017), Herrmann & Johnsrude (2020). As such, these models are linked to the classic model of attention and effort of Daniel Kahneman (Kahneman, 1973). For instance, in a 2016 consensus paper, listening effort was defined as ”the deliberate allocation of mental resources to overcome obstacles in goal pursuit when carrying out a task, with listening effort applying more specifically when tasks involve listening” (Pichora-Fuller et al., 2016). More recent work also analyzed the interaction of listening effort as defined in this way and affect, see Francis et al. (2016), Herrmann & Johnsrude (2020).

There is a large body of literature describing many different metrics of listening effort, usually categorized into self report (e.g., listening effort questionnaires), behavioral (e.g., reaction time, percent correct responses), and physiological measures (e.g., EEG based measures, pupil size, electrodermal activity, heart rate variability, hormonal measures). See, for example, Bernarding et al. (2010), Strauss et al. (2010), Mackersie & Cones (2011), Bernarding et al. (2013), Schafer et al. (2015), Seeman & Sims (2015), Francis et al. (2016), Bernarding et al. (2017), Bräannströom et al. (2018), Dimitrijevic et al. (2019), McGarrigle et al. (2021), Bräannströom et al. (2021), Giuliani et al. (2021), Francis et al. (2021), Fiedler et al. (2021), Paul et al. (2021), Ala et al. (2022), Haro et al. (2022) and Guijo & Cardoso (2018) for a broader literature review. There is, however, also a well described lack of correlation between these measures, which is generally attributed to the idea that different measures are sensitive to different dimensions of the complex construct that is listening effort (Strauss & Francis, 2017, Miles et al., 2017, Alhanbali et al., 2019, Lau et al., 2019, Colby & McMurray, 2021, Shields et al., 2023).

Therefore, in the context of potentially reorienting or shaping the pinna by means of auricular muscle activity, the differentiation between endogenous and exogenous factors for auditory stream selection (see Strauss et al. (2010), Strauss & Francis (2017)) is the main motivation behind the current study. Attempts to alter the physical properties of an auditory stimulus by changing the shape of the pinna and/or ear canal to aid auditory stream segregation seem to be a plausible function of the vestigial pinna–orienting system. Thus we hypothesize that such attempts should be reflected in the electromyogram of the auricular muscles.

Based on the findings from Strauss et al. (2020), we will focus our analysis on the SAM. We will remove confounding factors, such as the spatial separation between target and distractor streams (because this introduced an ipsi/contralateral lateralization effect). Furthermore, the demand dimension will be manipulated by two factors: the fundamental frequency differences between target and distractor, as well as by general signal-to-noise (SNR) changes, see the discussion in (Pichora-Fuller et al., 2016).

## 2 Material and Methods

### 2.1 Participants

20 participants were recruited for this study (12 male, 8 female; 18 right-handed, 2 left-handed). They were, on average, 28 *±* 4 years old and normal-hearing (according to a pure tone audiogram using test frequencies from 125 Hz to 8 kHz). The experiment was explained to every participant in detail before they signed a consent form. The study was approved by the responsible ethics committee (ethics commission at the Ärztekammer des Saarlandes, Saarbrücken, Germany; Identification number: 76/16)

### 2.2 Experimental Setup

Participants were seated in a chair in the center of a a 3*×*3*×*3 m cubicle made of heavy stage molton (900 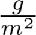) to reduce sound reflections. In order to avoid head movements during the experiment, the participants heads were placed on a chin rest. Two active loudspeakers (KH120A, Neumann, Germany) were placed in front (0°) and behind (180°) the participants, at a distance of 1.15 meters and at head level. A screen with a fixation cross was placed 80 cm away from the participants head,below the loudspeaker placed at 0°, therefore not blocking the loudspeaker. The setup was calibrated using a Brüel & Kjær Type 2250 Sound Level Meter. A stimulation PC controlled the loudspeakers via an external USB sound interface (Scarlett 18i20, Focusrite Plc., UK), generating audio and trigger signals at 44.1 kHz.

### 2.3 Stimuli and Tasks

Three different podcasts/audiobooks were used as target and distractor stimuli. For the target streams, segments of an audiobook, spoken by a female speaker, who briefly discusses a variety of topics (approximately 1 minute per topic) were used. Two different radio podcasts, one spoken by a male speaker, one by a female speaker (similar to the target) were utilized as distractors. Because their runtime was shorter than the experiment, we systematically shifted the starting points of the distractor podcasts so no trial had the same “background” noise. Pauses, defined as the 100 ms long moving average of the rectified waveform being less than 0.001, in all stimuli were removed (these parameters were determined experimentally) in order to increase the overall difficulty by increasing the information density and removing random unmasking effects.

**Figure 1:**
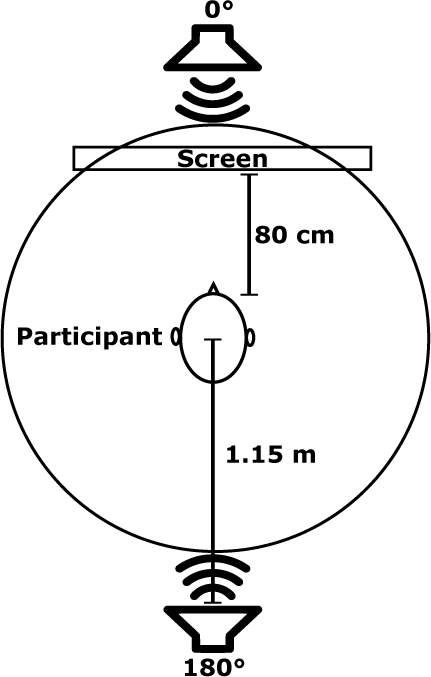
Experimental setup indicating the positions of the loudspeakers around the participant.

During the experiment, participants were instructed to attend to the target podcast, while ignoring the distractor podcasts. Target and distractors were always presented from the same loudspeaker (either from 0° or 180°), i.e., there were no spatial cues segregating target and distractor streams.

Three different levels of required listening effort (referred to as LE conditions) were generated, based on the target/distractor signal-to-noise-ratio (SNR), the number of distractors, as well as the distractor pitch. In the condition designed to require the least listening effort (low), the distractor was 10 dB softer then the target podcast (+10 dB SNR). Additionally, the distractor was a male speaker, meaning a high pitch difference between target and distractor. For the medium LE condition, an additional, female distractor was added with a pitch similar to the target podcast. Furthermore, both distractors combined were only 2 dB softer than the target (+2 dB SNR). For the high LE condition, the SNR was further lowered to -2 dB by increasing the distractor intensities. Table 1 summarizes the stimulus intensities of the three aforementioned LE conditions. Note that the target intensity was identical in all three conditions and only the distractor(s) varied.

**Table 1:** Stimulus intensities and SNR values for the conditions designed to require low, medium and high amounts of listening effort.

| LE condition | Target<br>[dB LAeq]<br>(female speaker) | Distractor 1<br>[dB LAeq]<br>(male speaker) | Distractor 2<br>[dB LAeq]<br>(female speaker) | Sum Distractors<br>[dB LAeq] | Sum All<br>[dB LAeq] | SNR [dB] |
| --- | --- | --- | --- | --- | --- | --- |
| Low | 50 | 40 | N/A | 40 | 50.4 | +10 |
| Medium | 50 | 45 | 45 | 48 | 52.1 | +2 |
| High | 50 | 49 | 49 | 52 | 54.1 | -2 |

It should be noted that we specifically avoided any spatial separation between target and distractor streams because strong non-spatial features, which we used to adjust the level of listening effort, are known to interact with EEG based spatial auditory attention measures, see Bonacci et al. (2020). Therefore, if the low, medium and high LE conditions used spatially separated target and distractor streams, the spatial component might be negligible in the low (Bonacci et al., 2020), but dominant in the high, LE condition (Fintor et al., 2022).

Considering two stimulus directions and three LE conditions, a total of six combinations were possible. For each combination, two trials of 5 minutes and 10 seconds each (12 trials in total) were recorded. In the first 5 seconds of each trial, only the target speaker was active, giving the participant the opportunity to solely focus on the target stream. During the next 5 seconds, the distractor(s) linearly faded in. Only the remaining 5 minutes of the trial, during which the distractor steams were at full intensity, were used for data analysis. The presentation order of the six combinations was randomized and balanced across participants, but the trials of each combination were always the same.

After each trial, participants rated the required listening effort on a 7-point scale (from effortless to extreme) and gave an approximate number of how often they lost the target stream during the trial (up to 10). Then, to ensure that participants had not given up during the experiment, they were instructed to recall the topics discussed in the corresponding target podcast trial (on average 4 topics per trial), as well as answer content related questions. It should be noted that listening effort scores and the number of target streams lost were always asked before the topic recall and content related questions in order to avoid any bias based on the participants’ impression of how well they were able recall the topics and answer the associated questions. Furthermore, participants were given the option to take breaks between trials.

### 2.4 EMG Data Acquisition

Passive Ag/AgCl electrodes were used to record EMG signals from the left and right superior auricular and postauricular muscles (SAM, PAM), as well as the masseter muscles (M. masseter). For the SAM, which is the largest extrinsic auricular muscle with an average width of 5 cm and a height of 4.7 cm (Chon et al., 2021), five electrodes per side were placed to cover the muscle, see Fig 2. This wide spread of electrodes is justified by the large variations of the size of the SAM: as reported by Chon et al. (2021), the central length ranges from 2.5 to 6 cm, and the width from 4 to 6.5 cm. Each PAM was recorded using electrodes placed directly on the PAM and the rear of the pinna (the position that yielded the largest PAM reflex in O’Beirne & Patuzzi (1999)). The M. masseter was recorded by placing one electrode on each side, slightly below the temporomandibular joint (the point which strongly protrudes during teeth clenching). One concern was that participants could, due to the long use of the chin rest, move their jaw or clench their teeth during the experiments, activating the temporal muscle which could be picked up by the electrodes placed on the SAM. Because both the M. masseter and temporalis muscle (M. temporalis) are involved in these movements, signals recorded from the M. masseter will later on be used to remove potentials artifacts. All electrodes were initially referenced against the ground electrode, which was placed at the upper forehead (Fpz). All signals were recorded at 4800 Hz using a commercially available, direct current (dc)-coupled, biosignal amplifier (g.USBamp, g.tec, Austria).

**Figure 2:**
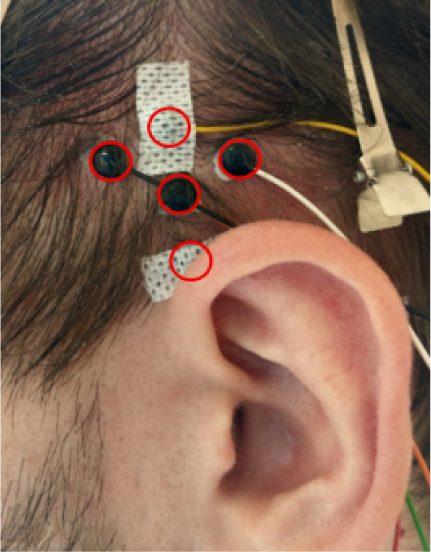
Positions of the five electrodes used to cover the SAM.

### 2.5 Signal Processing

Signal processing and statistical analyses were performed using Matlab 2020a (Mathworks, USA), IBM SPSS Statistics 28 (IBM Corp, USA), and R 4.2.1 (R Core Team, 2022). Raw EMG signals were initially re-referenced to bipolar signals: For the PAM, both electrodes on one side were used for re-referencing, resulting in one bipolar PAM signal per side. For the SAM, as there was no a priori knowledge which reference would be “ideal”, all 10 possible bipolar references per side were calculated. Signals from the left and right electrodes placed on the M. masseter were used to calculate one bipolar M. masseter signal. All signals were 10-500 Hz bandpass filtered (3rd order butterworth) and a 50 Hz IIR comb filter was applied (all filters were implemented as zero-phase filters). Signals were then segmented into 1 second long, non-overlapping segments. Next, artifact rejection was performed, independently for every trial and participant, based on two metrics. The first metric is based on the M. masseter signal. If the mean absolute value of any (1 second) segment exceeds 10 *µ*V, then the corresponding segments of the auricular signals were flagged as artifacts and discarded from further analysis (the threshold of 10 *µ*V for all participants was determined experimentally by instructing them to sit relaxed and then purposefully clench their teeth). The second metric was based on the auricular signals themselves. The energy of every 1 second long segment was calculated, and any segment that deviated by more than two standard deviations from the mean energy of the corresponding trial was removed from further analysis. Table 2 summarizes the artifact rates based on all participants and trials.

**Table 2:** Averaged artifact rates and standard deviations based on all partici-pants and trials.

| Muscle | Artifact Rate [%] |
| --- | --- |
| SAM Left | $5.7 \pm 3.6$ |
| SAM Right | $5.7 \pm 3.5$ |
| PAM Left | $5.9 \pm 4$ |
| PAM Right | $6 \pm 3.8$ |
| M. Masseter | $3.2 \pm 4.2$ |

Because, as mentioned above, there was no a priori assumption regarding the best SAM reference, the next step was to survey all possible references and check for apparent trends. A very interesting observation was that on both left and right SAM, contrasts between LE conditions became, on average, larger approximately 150 seconds into the trial and tapered off in the last few seconds. This observation motivated us to focus on the time frame from 150 to 294 seconds of every trial. Furthermore, results from different references appeared to be very similar, meaning an objective performance metric had to be implemented. We decided on a metric, similar to the F-statistic used in the ANOVA, which maximizes differences between classes (in this case, levels of required listening effort), but also takes into account the variance within each class.

For this purpose, the non-normalized segmented energy signals of all four trials (from one participant and reference) corresponding to one LE condition were concatenated and will be referred to as *y_i,j_*, where *i* indicates the LE condition and *j* a 1 second segment. 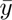*_i_* will be the corresponding group average of the *i*-th LE condition across all *j* valid segments. 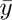 is the grand average of all *N* valid segments across all *k* LE conditions. A distance metric can then be calculated as follows:

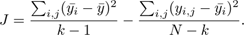

It is important to note that this metric is not biased towards a specific LE condition. This metric was calculated independently for every participant and left/right SAM and was used to determine the ideal reference per participant. Finally, for every direction (0° and 180°) and all three LE conditions, the mean energy from all valid 1 second long segments was calculated. These averaged values were then z-normalized, independently for every participant and muscle (left/right PAM, SAM, M. Masseter), and subjected to statistical analysis by means of a two-way repeated measures ANOVA (2 directions *×* 3 LE conditions).

## 3 Results

Figure 3 displays the averaged subjective ratings (listening effort and how often the participants lost the target stream) per LE condition after z-normalization, as well as the scores of the correctly answered questions and topics recalled. Re-peated measures ANOVAs with LE condition and stimulus direction as factors indicated significant effects of the LE condition for the subjective listening effort rating (*F* (2, 38) = 336.332*, p <* 2 *·* 10*^−^*^16^) and how often participants lost the target stream (*F* (2, 38) = 303.929*, p* = 2.1 *·* 10*^−^*^15^). Pairwise t-tests (*df* = 19, Bonferroni corrected) show that, for self reported listening effort and number of target streams lost, each LE condition significantly differs from another. The averaged ratings of the listening effort scale almost resemble a straight line, whereas for the target lost metric, the difference between the low and medium LE condition is (on average) much smaller than the difference towards the high LE condition. Non-normalized, averaged values for the number of target streams lost were 1.038 *±* 1.169, 1.95 *±* 2.028 and 5.919 *±* 2.748, respectively. Supplementary Figures 8 and 9 split these data further into all six stimulus direction and LE condition combinations, as well as according to the presentation order to check for any presentation order effects. When split into the six combinations, the subjective data display the same pattern, regardless of stimulus direction (0° and 180°), and there is no apparent rise or fall when the data is organized according to the presentation order. Regarding the question scores, significant main effects were observed for both LE condition and stimulus direction(LE condition: *F* (2, 38) = 6.696*, p* = 0.003, stimulus direction: *F* (1, 19) = 11.715*, p* = 0.003). Participants made significantly more errors in the high LE condition, compared to the low LE condition (mean percent correct scores for low, medium, and high: 82.35%, 72.08%, 63.23%). As for recalling the topics, there was only a significant main effect of LE condition (*F* (2, 38) = 11.637*, p <* 0.001). Significantly fewer topics were recalled during the medium LE condition compared to either the low or high LE condition (mean values for low, medium and high: 82.21%, 73.26%, 86.44%).

**Figure 3:**
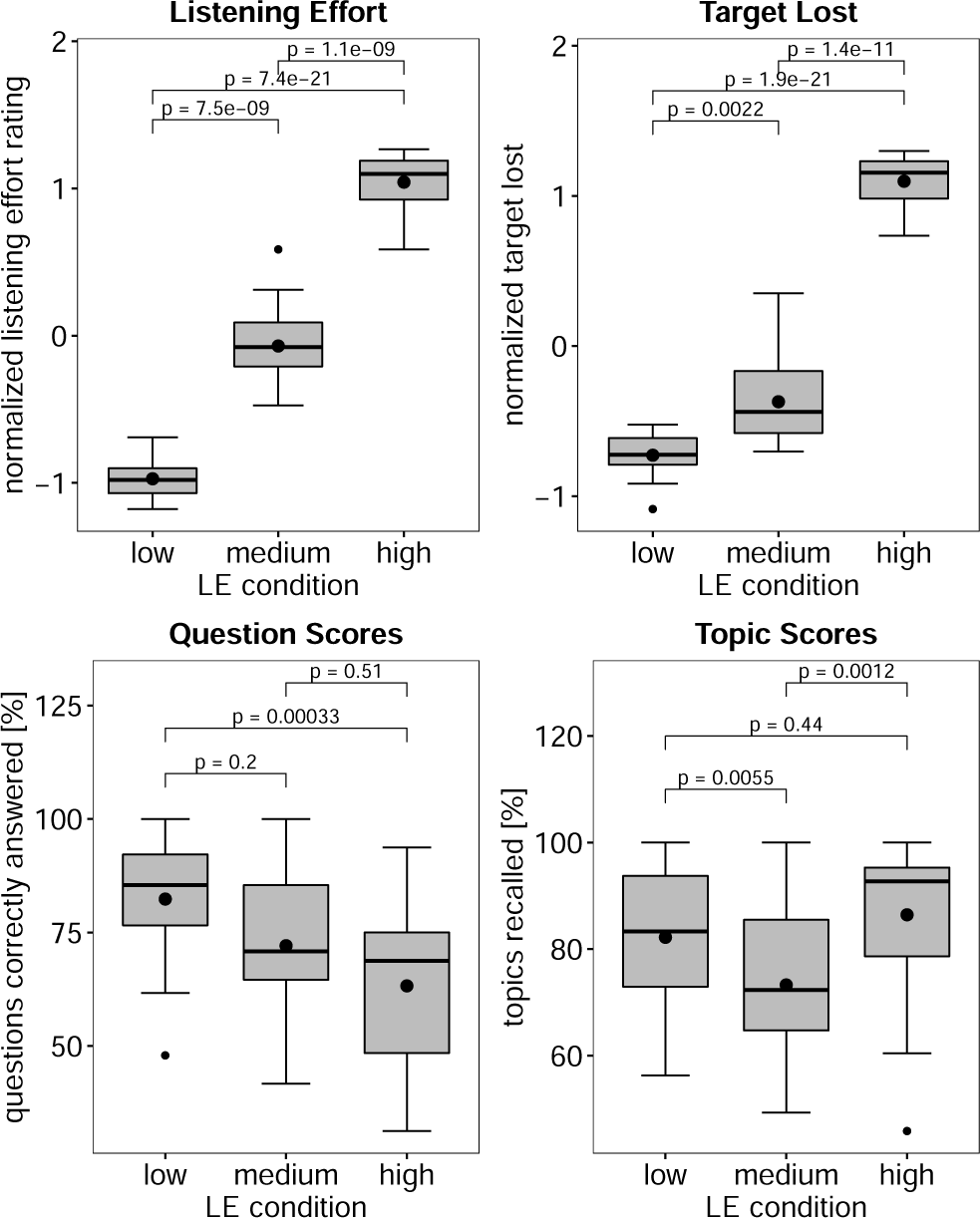
Averaged values of the normalized listening effort and target lost ratings, as well as percentages of correctly answered questions and topic recall scores. P-values were obtained using Bonferroni corrected paired t-tests (*df* = 19).

Figure 4 shows the normalized and time-resolved plots of the left and right SAM, averaged across all trials and participants, in 10-second steps, to generate a more smoothed curve. Note that only data from the second half (seconds 150- 294) were used for analysis, as the contrast between conditions is much smaller in the first half.

**Figure 4:**
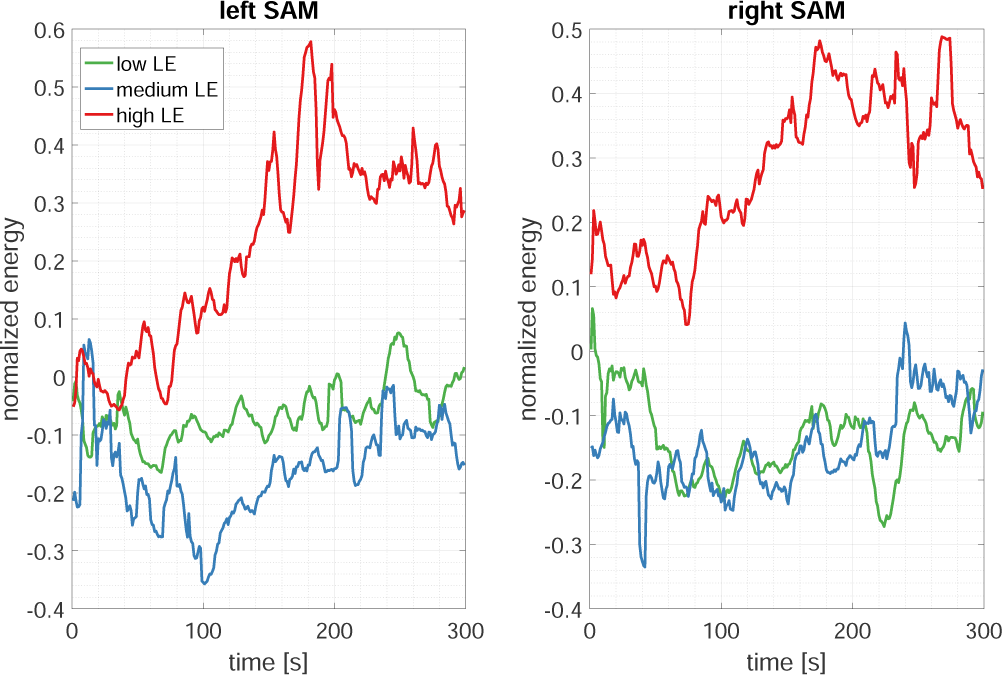
Time-resolved normalized activity of the left and right SAM depending on the three LE conditions. On both SAMs, the difference between the high and the low/medium LE conditions is, on average, largest in the second half (seconds 150-294) of the trials.

For both left and right SAM, two-way repeated measures ANOVAs revealed no significant interaction between stimulus direction and LE conditions as well as no significant main effect of direction. However, the LE condition had a statistically significant main effect on the outcome of both left (*F* (2, 38) = 7.594*, p* = 0.002) and right (*F* (2, 38) = 7.281*, p* = 0.002) SAM. Pairwise comparisons (paired t-tests, Bonferroni corrected) show that for both SAMs, results from the high LE condition are significantly larger than the low (left SAM: *t*(19) = 3.128*, p* = 0.017; right SAM: *t*(19) = 3.345*, p* = 0.01) and medium (left SAM: *t*(19) = 3.262*, p* = 0.012; right SAM: *t*(19) = 2.737*, p* = 0.039) LE condition, whereas differences between low and medium were not significant, see Figure 5. Supplementary Figure 10 shows the data from both SAMs split into all six combinations and supplementary Figure 11 demonstrates that there is no obvious effect of presentation order.

**Figure 5:**
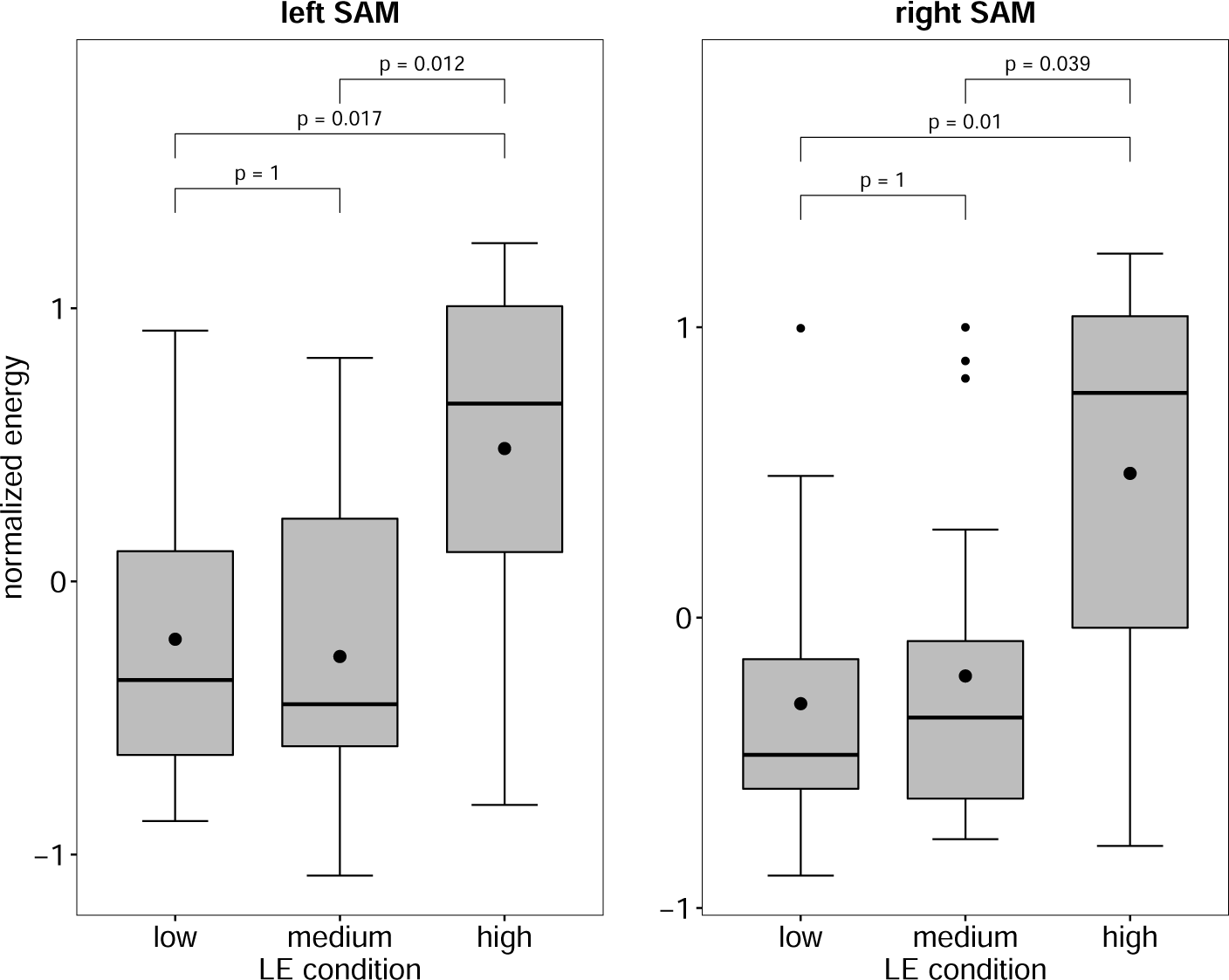
Boxplots of the normalized energy of the left and right SAM, depending on the LE condition. Black dots indicate the arithmetic means. P-values were obtained using Bonferroni corrected paired t-tests (*df* = 19).

Considering the left and right PAM, as well as the M. Masseter, no significant interaction between stimulus direction and LE condition, nor any main effect of LE condition was observed, see Figure 6. While there was also no main effect of stimulus direction in the data recorded from the M. Masseter, both left and right PAM displayed a significant effect of stimulus direction (left PAM: *F* (1, 19) = 12.806*, p* = 0.002; right PAM: *F* (1, 19) = 10.943*, p* = 0.004). It should be noted that in both cases, the PAM signals recorded while the par-ticipants were focusing on the back loudspeaker (180°) were significantly larger than when the audio streams were were presented from the front (0°), see Figure 7. Similar to the results from the SAM, data split into all six combinations as well as rearranged into the presentation order are displayed in supplementary Figures 12 and 13.

**Figure 6:**
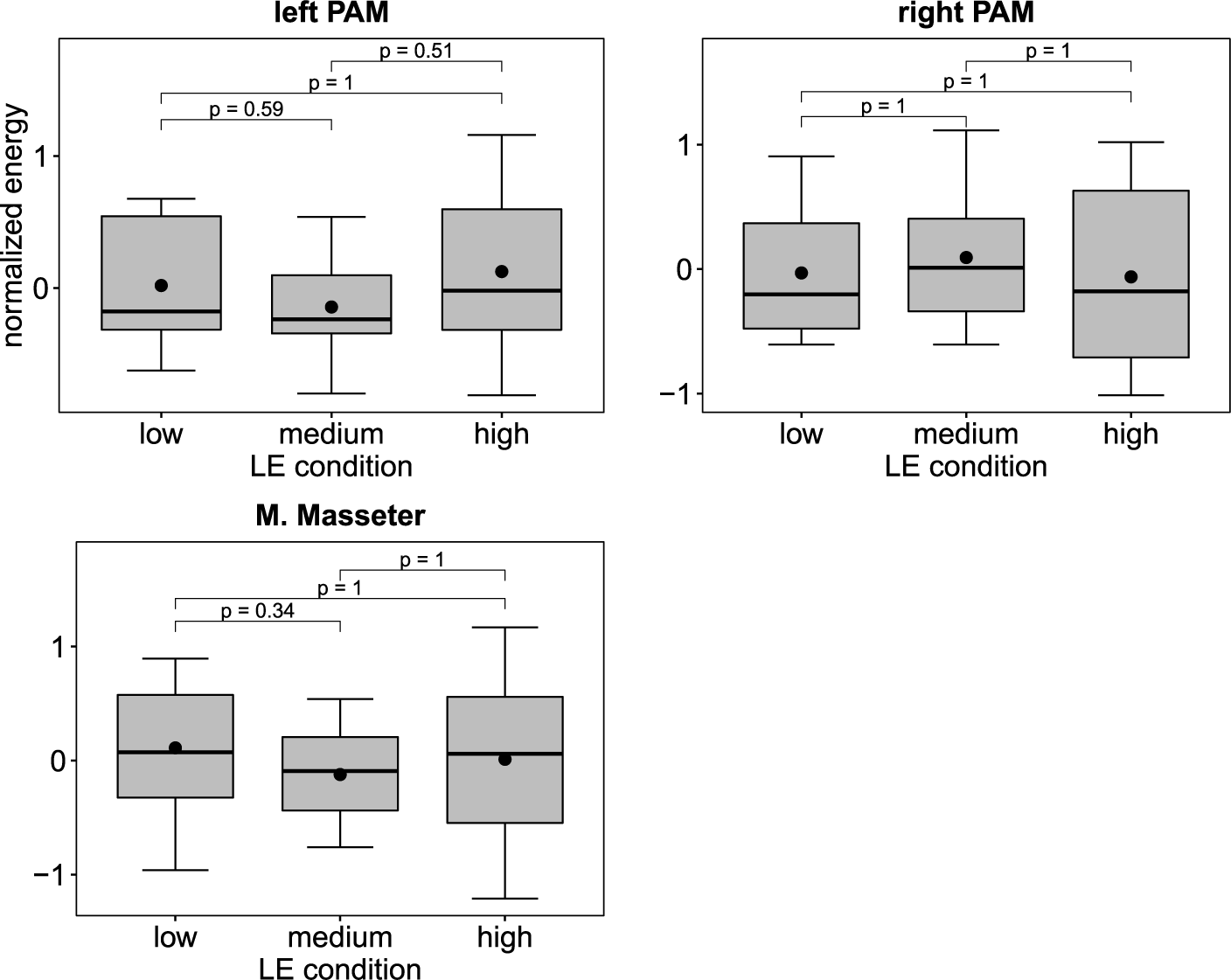
Boxplots of the normalized energy of the left and right PAM and the M. Masseter, depending on the LE condition. P-values were obtained using Bonferroni corrected paired t-tests (*df* = 19).

**Figure 7:**
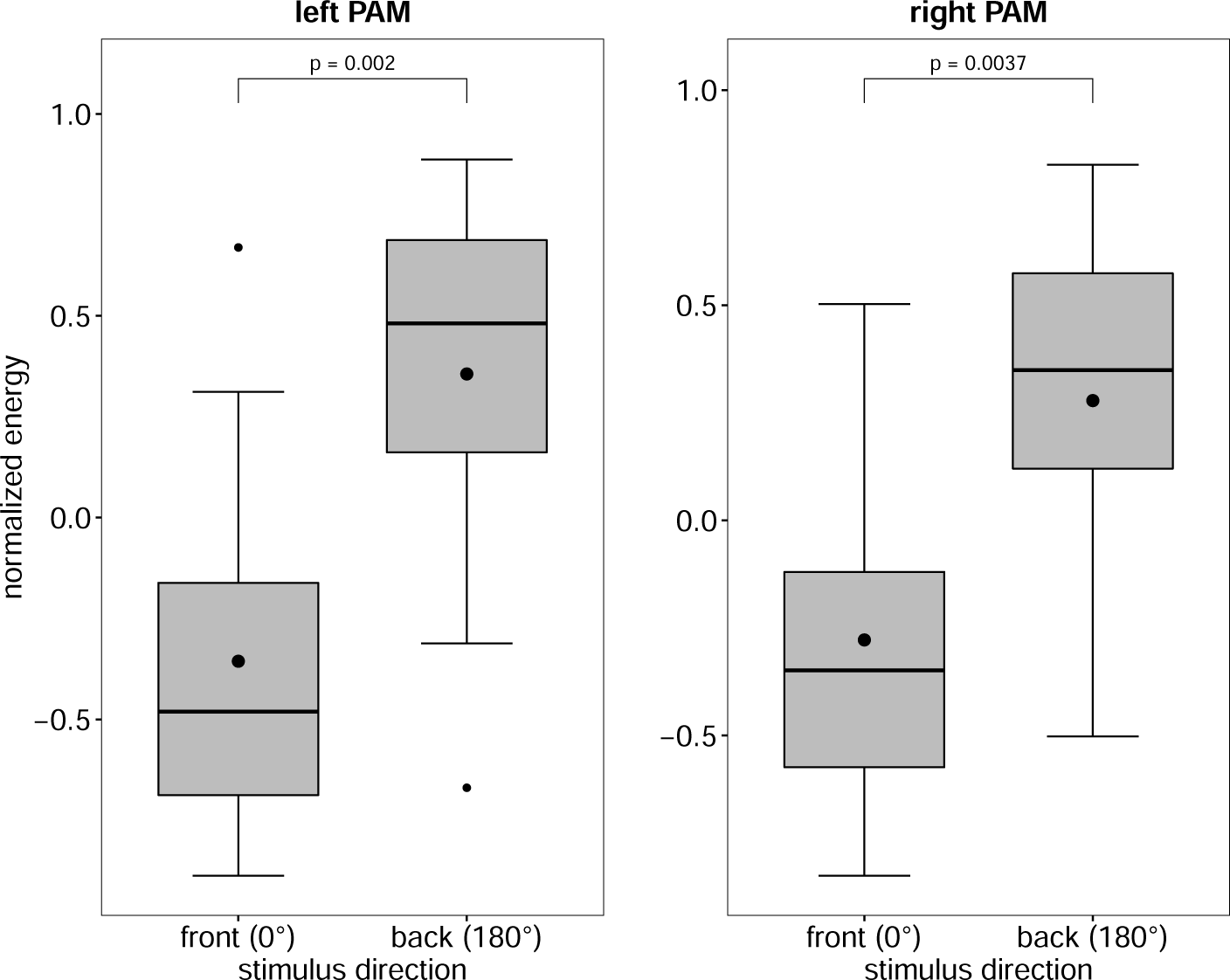
Boxplots of the normalized energy of the left and right PAM, depending on the stimulus direction. P-values were obtained using Bonferroni corrected paired t-tests (*df* = 19).

## 4 Discussion

Sustained activity of auricular muscles has been shown to reflect the spatial direction of auditory attention (Strauss et al., 2020), using a vestigial pinna-orienting system (Hackley, 2015). Based on these findings, we designed an experiment to check if this vestigial system could also be active during more generalized scenarios involving effortful listening. We generated conditions that require several distinct levels of LE, based on the number and pitch of distractors (Pichora-Fuller et al., 2016), while purposefully not spatially separating target and distractor streams to avoid lateralization effects (as reported in Strauss et al. (2020)). At the same time, we included two levels of stimulus direction (presentation of all streams from either 0° or 180°), because auricular responses were reported to be larger when stimuli were presented from outside the participants’ field-of-view (Strauss et al., 2020).

We found that signals from both left and right SAMs displayed significantly more activity during the high LE condition, compared to both low and medium. Differences between low and medium LE conditions were, however, not significant. A surprising finding was that the EMG activity of both SAMs generally increased over time and plateaued towards the end. It is surprising insofar as that the sustained SAM activity in response to spatial attention reported in Strauss et al. (2020) displayed a declining trend or remained stable over time, which might be attributed to the detrimental effect of prolonged time on task as described in (Sarter et al., 2006). In the design phase of the study, we specifically decided to record shorter, and therefore, more trials (2 *×* 5 minutes instead of one 10 minute long trial), because we initially speculated that the EMG activity could display a downwards trend and additionally wanted to avoid participants disengaging from the task due to fatigue or demotivation, which plays a pivotal role in listening effort research (Herrmann & Johnsrude, 2020, Francis & Love, 2020). The implication for future studies regarding listening effort using auricular muscles is therefore that if trials are too short, they may fail to capture this effect. Conversely, it would be interesting to study the time course of the SAM beyond the 5 minute mark in order to assess how long this effect lasts, and if we actually captured the “maximum”.

A similar pattern using EMG recorded from the frontalis muscles for three different levels of task demand was reported by Mackersie & Cones (2011), where EMG activity significantly increased from medium to high demand only. However, Mackersie & Cones (2011) recorded different muscles and utilized different stimuli/paradigms than the present study, making a direct comparison difficult. Francis et al. (2021) recorded EMG from the corrugator supercilii (which is very close to the frontalis muscle), but was unable to observe any significant effect between two different levels of LE. The authors speculate that this could in part be due to the low affective valence of the stimuli used, which the corrugator supercilii is an indicator of. To our knowledge, there is currently no evidence to suggest an effect of affective valence of auditory stimuli on the SAM response. Therefore, given that the stimulus material across LE conditions was from the same audiobook and speaker, the affective valence of the stimuli should be mostly constant throughout the experiment and should not be a confounding factor.

Interestingly, Francis et al. (2021) furthermore report an effect of SNR on self-rated effort, but not on physiological measures, which includes the corrugator supercilii. They suggest that within a certain stimulation range, sound-level related effects are negligible on physiological responses. This interpretation could be supported with the data of the present study: the sound-level differences between the low and medium LE condition are large enough for significant differences in self-reported LE, but not for the physiological response (in this case the SAM). The high LE condition, on the other hand, might just cross a threshold for which some physiological measures become sensitive for.

Comparing the collected SAM data to the self reported listening effort scale, the SAM does not capture the difference in the reported listening effort ratings between the low and medium LE conditions, which is comparable to the difference between the medium and high LE condition. Perhaps this difference may be explained by a recent review by Shields et al. (2023), which analyzed the correlation between different measures of listening effort and concluded that correlation between effort questionnaires and physiological measures were mostly poor to fair, and only 28.8% were significantly correlated.

A different self reported measure (the number of how often participants lost the target stream), appears to capture the results of the SAM more closely, namely a much smaller difference between the low and medium LE condition. In Pichora-Fuller et al. (2016), the authors developed a three dimensional model, in which effort is a nonlinear function of demand and motivation. Assuming a constant level of motivation, the results from both the SAM and the target lost metric could easily fit onto such a curve: even if the demand between the low, medium and high LE conditions (as quantified by the self reported listening effort questionnaire) would be evenly spaced, the proposed non-linear relation between demand and effort could result in a negligible difference between low and medium LE conditions and substantial increase in effort during the high LE condition, see also the computational model of the demanded and exerted effort relation in Schneider et al. (2019). However, we did not ask *how long* participants lost the stream, which could potentially differentiate between the low and medium LE conditions. On the other hand, instructing the participants to keep track of the number and duration of how often they lost the target stream would be an additional task and might severely distract from their primary objective. Nevertheless, there is a growing consensus that measures of listening effort (or effortful listening) depend on different underlying dimensions,are not interchangeable, and depend on a complex interaction between external and internal factors, such as fatigue, motivation, and attention (McGarrigle et al., 2014, Strauss & Francis, 2017, Alhanbali et al., 2019, Herrmann & Johnsrude, 2020, Francis & Love, 2020). Therefore, we could interpret the recorded SAM data in the context of losing the target stream, which, on average, participants did once in the low, twice in the medium, and six times in the high LE condition. The vestigial pinna-orienting system could try to change the spectral properties of the pinna or the ear canal. This could lower the external (or perceptual) LE (as opposed to internal or cognitive LE, see Strauss & Francis (2017), Francis & Love (2020)) and therefore aid to “locate” the target stream.

Strauss et al. (2020) has shown that transient and sustained involuntary activity of auricular muscles can lead to visible movements or deformations of the pinna shape. If large enough, such movements might impact the head related-transfer function (Stitt & Katz, 2021). Whether or not movements of the auricular muscles can affect the shape of the pinna to such a large degree that a utilizable change is generated would require a dedicated, future study that includes appropriate video recordings. If attention-driven auricular movements are purely vestigial in our own species, clues as to their original function might be discerned from other primates. Directly stimulating the VIIth nerve branch to SAM in an anesthetized macaque, Waller et al. (2008) (see supplement) showed that maximum contraction yields an upward, essentially rigid, translation of the pinna relative to the ear canal. The pinna as a whole does not appreciably rotate or deform. A consequent shift in distance of the upper and lower walls of the concha (which is of special importance in determining the HRTF, see Stitt & Katz (2021)) might generate a simple, predictable change in the spectral properties of the proximal stimulus while maximizing the aperture of the ear canal. By contrast, isolated contraction of the PAM or AAM homologs yields more complex, multidimensional movements in which the tragus can occlude the ear canal.

The behavioral responses (question and topic recall scores) show a less clear picture. While the question scores do display a decline with increased required listening effort, only the difference between the low and high LE condition reached statistical significance. Topic recall scores even show significantly lower scores in the medium LE condition compared to both low and high. However, we should mention again that both scores were designed to check general participant compliance, i.e., whether participants stopped solving the task due to, for example, boredom (if scores in the low LE condition were very low), or if they gave up (low scores in the high LE condition). Both cases would could have effects on the physiological measures (Herrmann & Johnsrude, 2020). Note that trials had a varying number of topics (2-7) and associated questions (1-4) and were fixed to a corresponding LE condition. For example, a topic about animal behavior contained a question that many participants failed to answer and was always part of the medium LE condition. Interpreting these scores is therefore difficult, because there may be systematic bias present. This could also be an explanation for the significant effect of stimulus direction on the question scores. It is possible that the questions associated with the stimulus material presented form 180° are simply significantly easier. Nevertheless, both scores were, on average, above 63% (questions) and 73% (topics), which we believe indicates that participants consistently attempted to solve the paradigm and retained a certain level of motivation.

During post-hoc analysis, we observed that the activity of both PAM muscles was only significantly affected by the direction of the stimuli (0° vs. 180°), and not by the different LE conditions Specifically, PAM activity when attending audio streams from the back was significantly larger than attending the front.

While PAM activity was also larger when participants attended stimuli from the back in Strauss et al. (2020), their experiments focused specifically on spatial auditory attention, i.e., target and distractor streams were spatially separated. Furthermore, the loudspeakers in Strauss et al. (2020) were not placed directly in front of or behind the participants, but off-center at *±*30° and *±*120°. Combining the current results with Strauss et al. (2020), we can conclude that the PAM is generally more responsive to audio streams that are outside of the participants’ field-of-view. This could lead us to hypothesize that if the eye gaze cannot shift towards a stimulus, the vestigial pinna–orienting may activate the PAM to enhance the participant’s ability to focus on these sounds.

The primary potential confounding factor in this study was cross-talk from the M. temporalis, which is situated extremely close to the SAM. Specifically, we were concerned that participants might begin to grind their teeth during the experiment, due to their positioning on the chin-rest becoming uncomfortable over time, or as a general response to stress. The masseter muscle, which works in conjunction with the M. temporalis during mastication, should provide a good proxy signal to assess possible cross-talk between the SAM and M. temporalis. Because analysis of the M. masseter revealed no significant effect of LE conditions or stimulus direction, it seems unlikely that the signals recorded from the SAM are the result of cross-talk from the M. temporalis.

## 5 Conclusion

This study provides evidence that SAM activity can be an indicator for increased levels of listening effort. Unlike other reactions of the autonomic nervous system (e.g., skin conductance, pupil diameter, etc., see Mackersie & Cones (2011)), an increased activity of the vestigial pinna-orienting system (Hackley, 2015) could be interpreted as an attempt to alter the shape of the pinna or ear canal. This manipulation could potentially influence stimulus related factors in models of listening effort, such as the transmission factors as described in Pichora-Fuller et al. (2016), or external/exogenous factors in Strauss et al. (2010) and Strauss & Francis (2017). While increased activity of auricular muscles in response to automatic and intentional attention can lead to visible movements of the pinna (Strauss et al., 2020), it is currently not known if they are strong enough to achieve an actual benefit. Especially in the current experimental setup, without any spatial separation between target and distractor, orienting the pinna would be futile (unlike the spatial attention paradigm in Strauss et al. (2020)), which is perhaps why the effect was symmetrical in both SAMs and not lateralized. Nevertheless, future studies should focus on exploring the auricular muscles in the context of the multi-dimensional concept of listening effort (e.g., Alhanbali et al. (2019), Shields et al. (2023)), which was present in the current study when comparing the SAM results to the self reported listening effort ratings. In this context, focusing on the participants losing the target stream would be of interest, as this self reported measure seemed to resemble the SAM closer.

## Acknowledgments

We thank Christine Welsch for assistance with data collection.

## Data Availability

Anonymized data that support the findings of this study are available upon reasonable request from the authors, but will be uploaded to a public repository upon publication.

## Ethical Approval

This study was conducted in accordance with the principles embodied in the Declaration of Helsinki and in accordance with local statutory requirements. Participants signed a consent form after a detailed explanation of the experiment. The study was approved by the responsible ethics committee (ethics commission at the Ärztekammer des Saarlandes, Saarbrücken, Germany; Iden-tification Number: 76/16).

## A Supplementary Material

**Figure 8:**
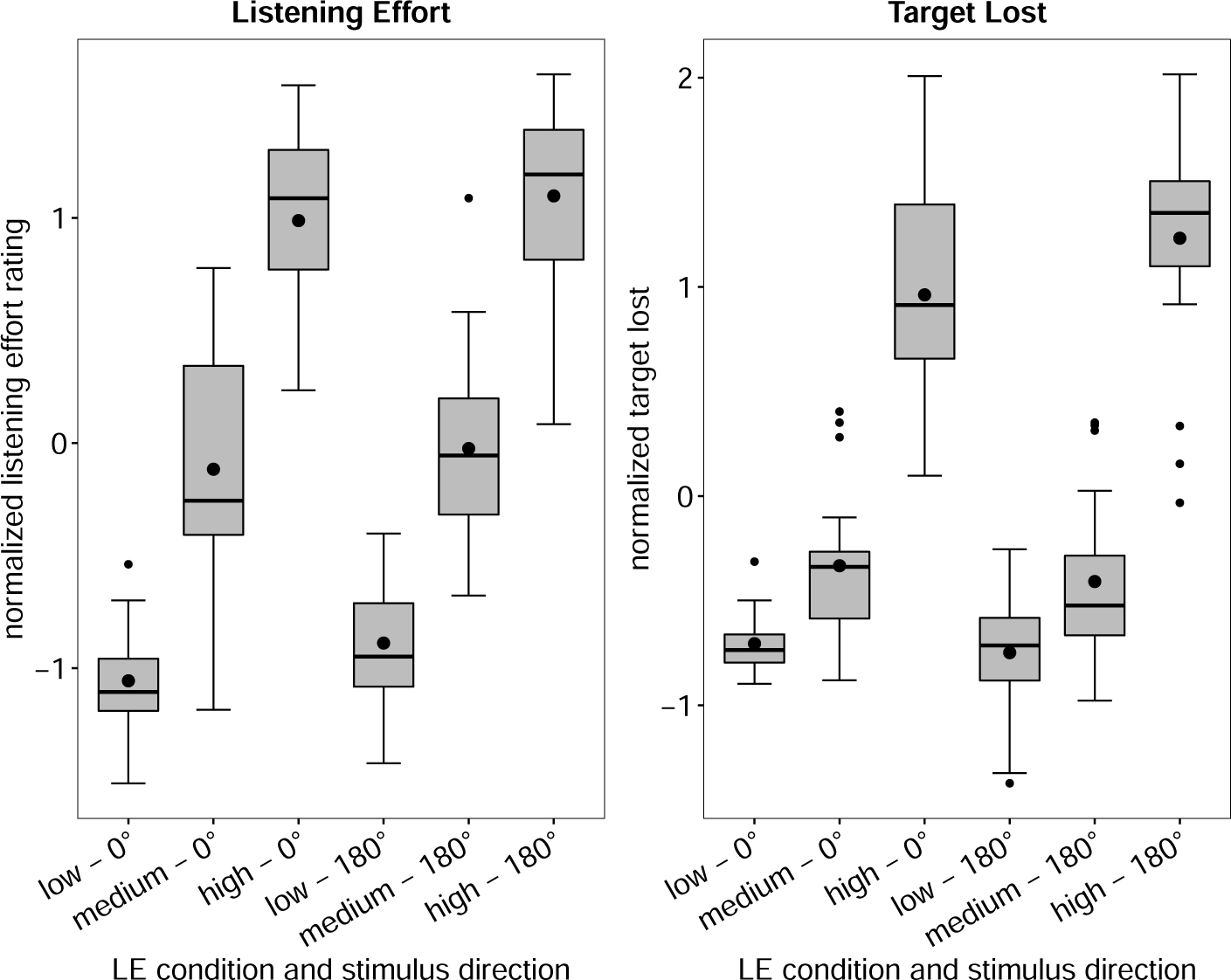
Boxplots of the normalized listening effort and target lost ratings, depending on the LE condition and stimulus direction.

**Figure 9:**
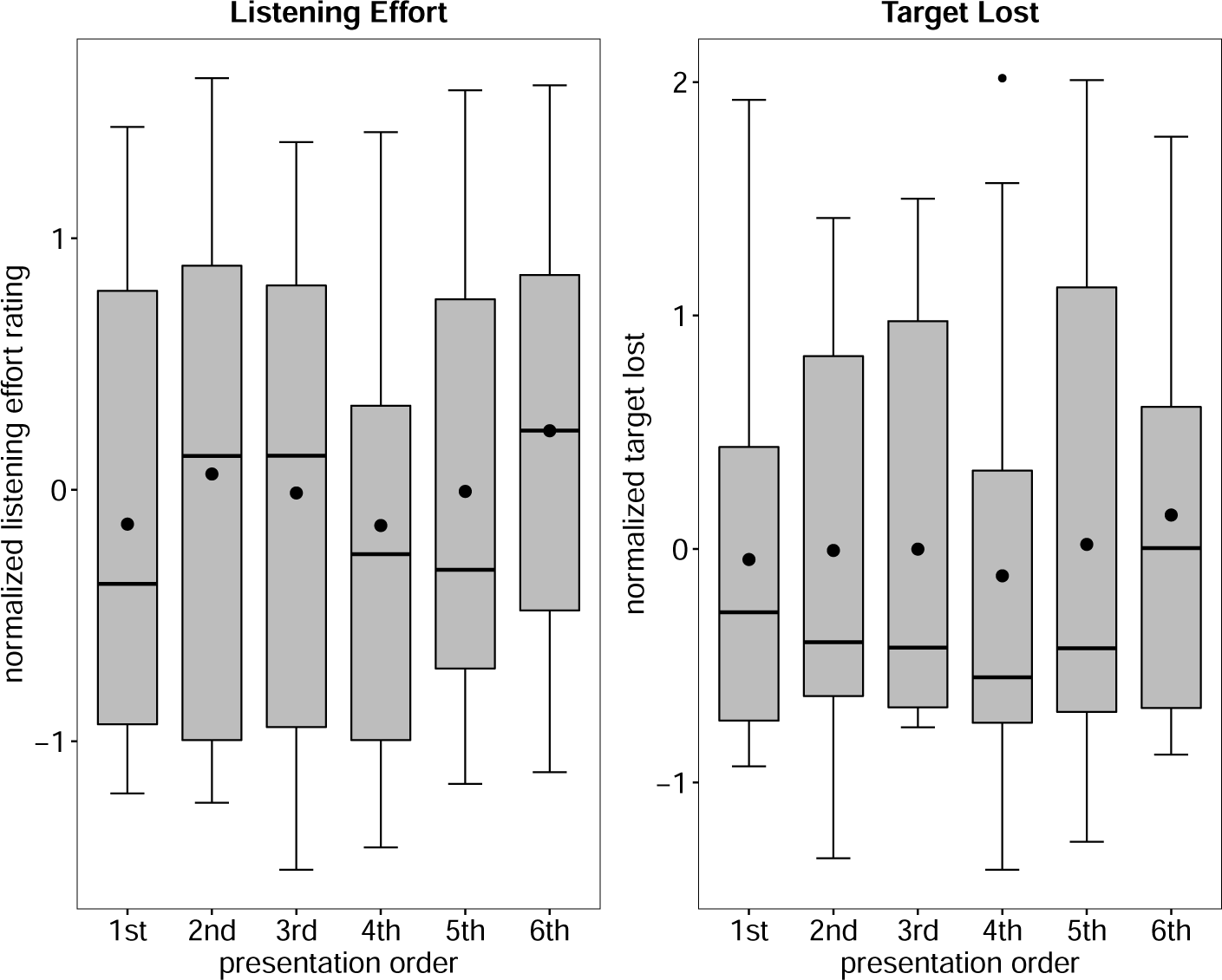
Boxplots of the normalized listening effort and target lost rating, depending on the presentation order.

**Figure 10:**
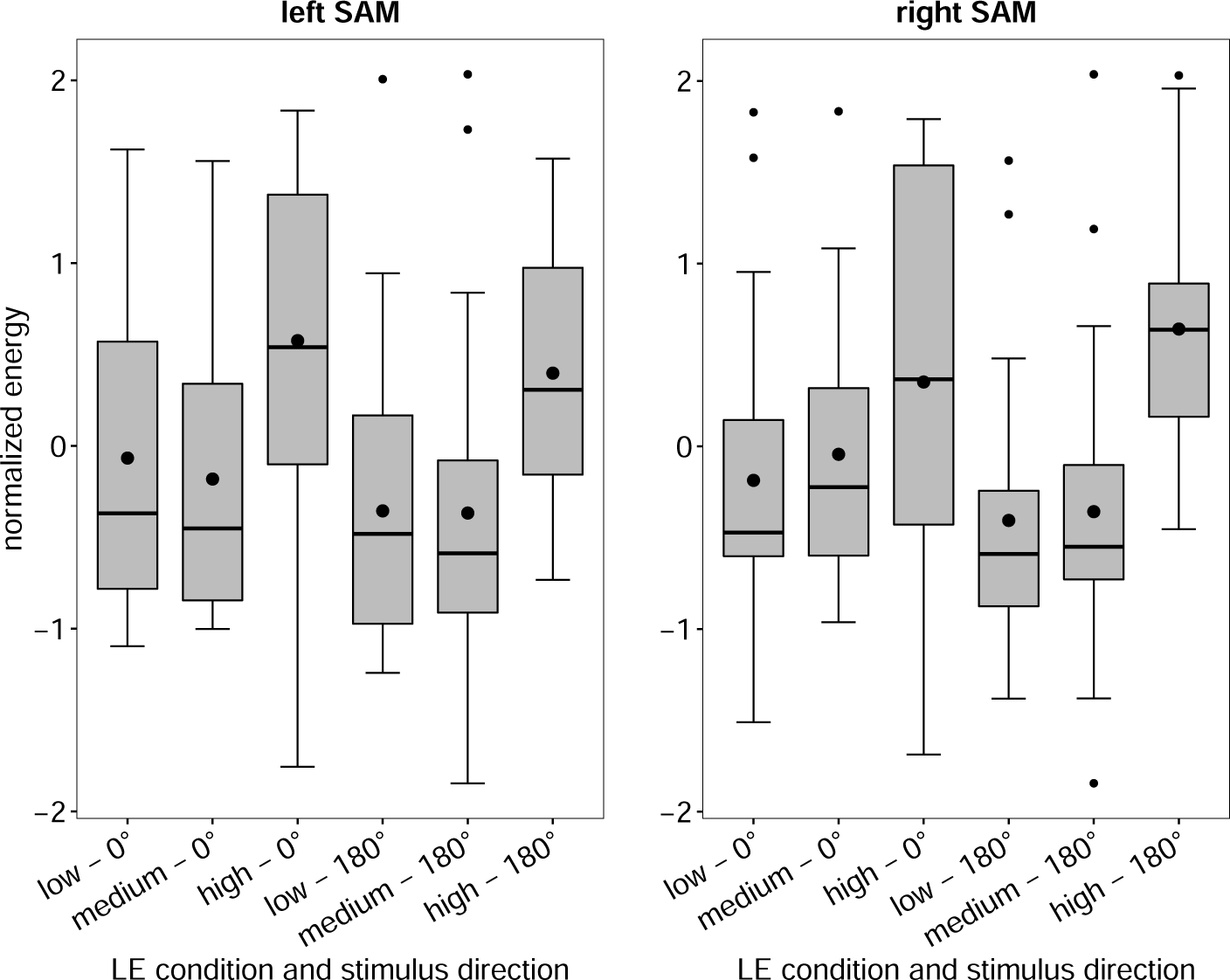
Boxplots of the normalized energy of the left and right SAM, depending on the LE condition and stimulus direction.

**Figure 11:**
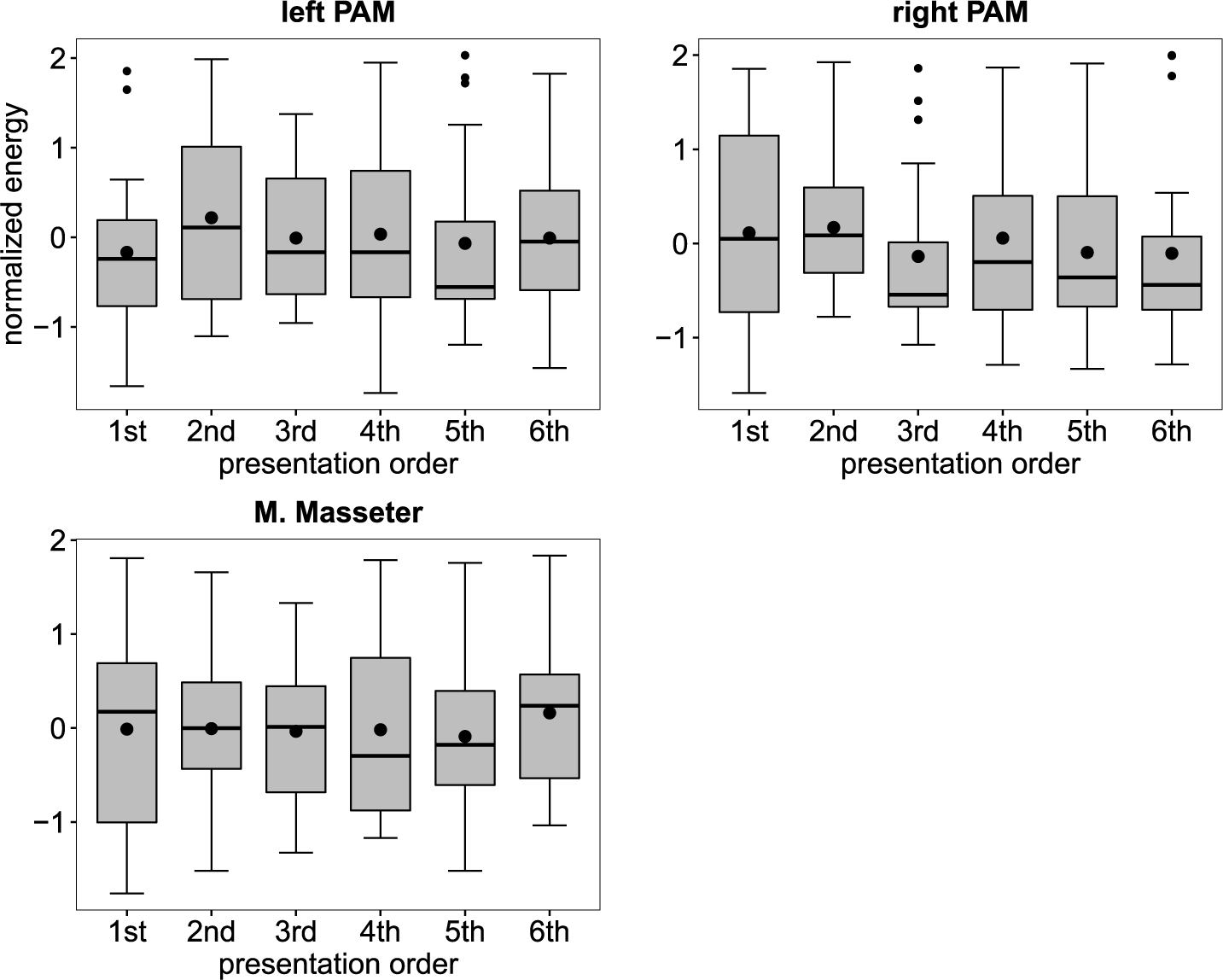
Boxplots of the normalized energy of the left and right SAM, depending on the presentation order.

**Figure 12:**
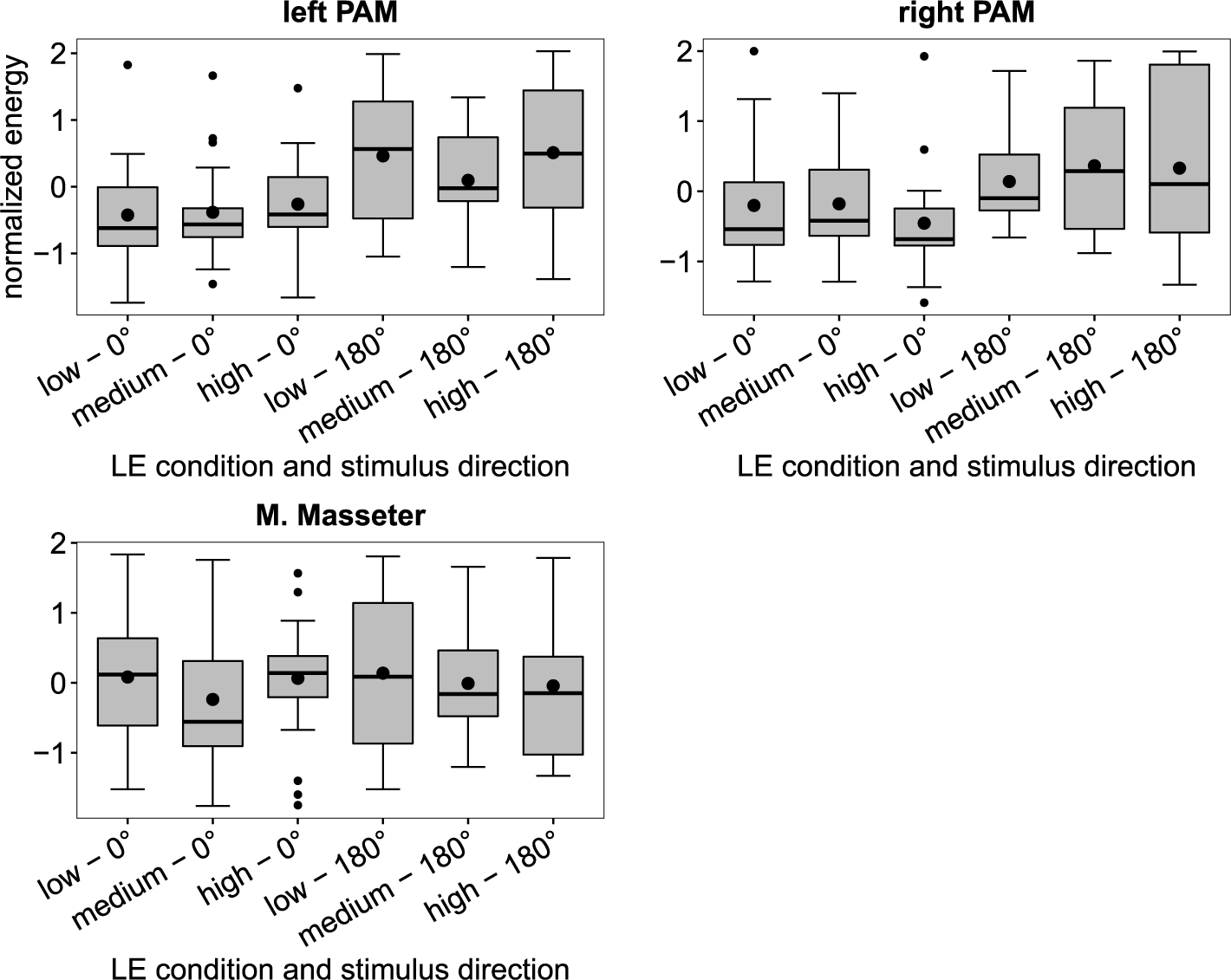
Boxplots of the normalized energy of the left and right PAM and the M. Masseter, depending on the LE condition and stimulus direction.

**Figure 13:**
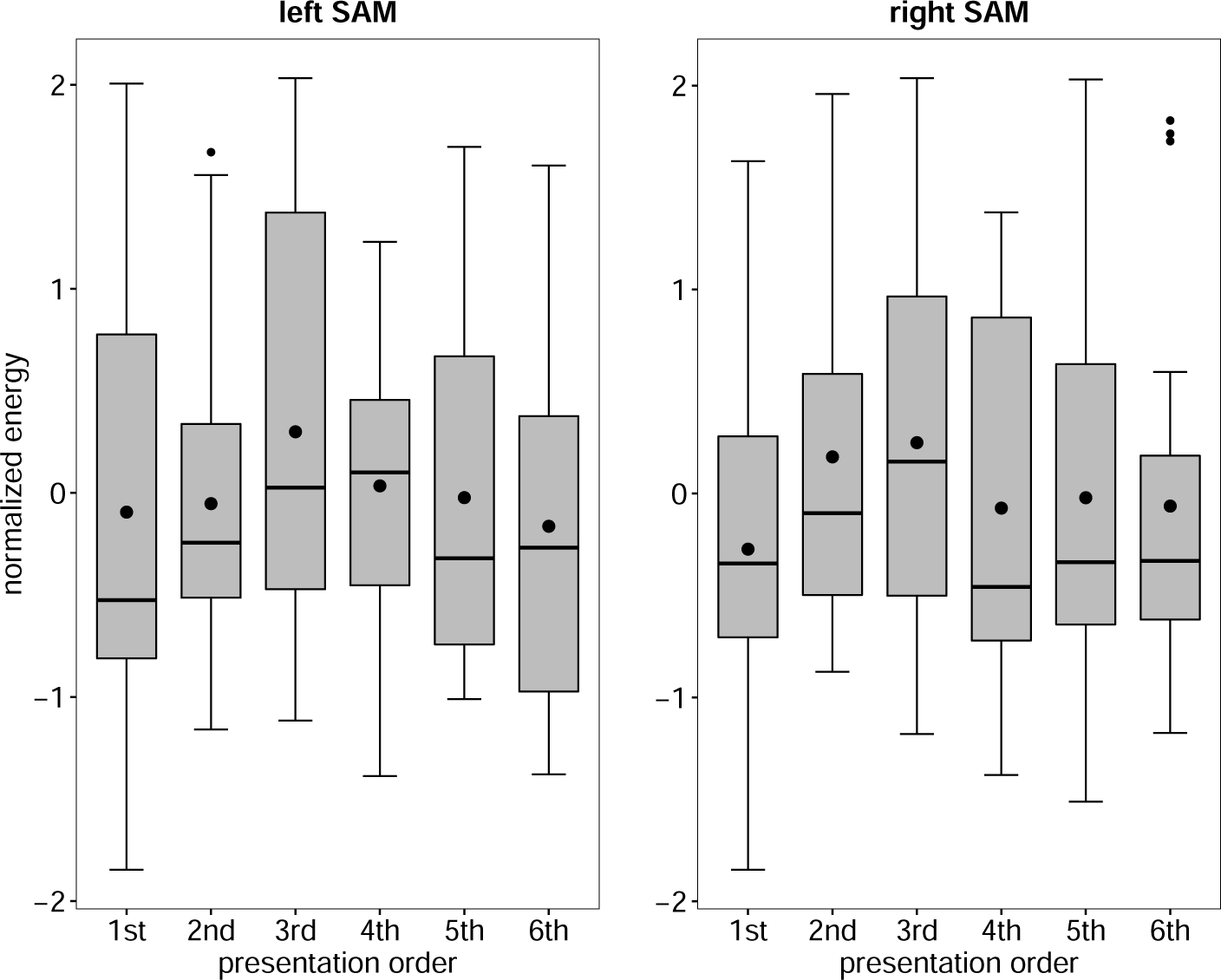
Boxplots of the normalized energy of the left and right PAM and the M. Masseter, depending on the presentation order.

